# Investigation of cell mechanics and migration on DDR2-expressing neuroblastoma cell line

**DOI:** 10.1101/2024.08.15.607761

**Authors:** Theadora Vessella, Esteban Rozen, Jason Shohet, Qi Wen, Hong Susan Zhou

## Abstract

Neuroblastoma is a devastating disease accounting for ~15% of all childhood cancer deaths. Collagen content and fiber association within the tumor stroma influence tumor progression and metastasis. High expression levels of collagen receptor kinase, Discoidin domain receptor II (DDR2), are associated with poor survival of neuroblastoma patients. Additionally, cancer cells generate and sustain mechanical forces within their enviroment as a part of their normal physiology. Despite this, whether collagen activated DDR2 signaling dysregulate these migration forces is still elusive. To address these questions, a shRNA DDR2 knockdown neuroblastoma cell line (SH-SY5Y) was engineered to evaluate the consequence of DDR2 on cellular mechanics. Atomic force microscopy and traction force microscopy were utlizing to unveil the biophysical altercations. DDR2 down-regulation was found to significantly reduce proliferation, cell stiffness, and cellular elongation. Aditionally, DDR2 down-regulated cells had decreased traction forces when plated on collagen coated elastic substrates. Together, these results highlight the important role that DDR2 has in reducing migration mechanics in neuroblastoma and might be a promising target for future therapies.

## 1. Introduction

Metastasis is a hallmark of aggressive cancer, and the metastatic cascade is characterized by migrating cancer cells interacting with their local microenvironment to sense biomechanical properties and adapt their migratory phenotype. Neuroblastoma (NB) is the most common extracranial solid tumor in childhood, with metastatic spread seen in half of the patients [1, 2]. NB accounts for ~8% of all childhood cancers, but around 15% of pediatric cancer mortality [3]. Despite efforts to treat the cancer, 50-60% of high-risk patients will relapse within two years of a diagnosis [2]. Understanding how the extracellular matrix (ECM) regulates various cellular functions such as cell spreading, migration, proliferation, and differentiation [4–6] is vital to improve the therapeutics for NB.

Collagen is the major component of the extracellular matrix (ECM) in most mammalian tissues [7]. Cells bind to the collagen through adhesion receptors, these receptors then activate actomyosin machinery that is capable of generating forces required for migration and delivering important signals for growth and survival [8]. In addition to integrins, the discoidin domain receptors (DDR), DDR1 and DDR2, are well-known collagen receptors. The DDRs are widely expressed in human and mouse tissues with distinct distributions; DDR1 is enriched in epithelial cells and DDR2 is mainly expressed in mesenchymal cells [9]. Compared to DDR1, DDR2 has a greater specificity for fibril-forming collagen [10–12]. While DDR2 plays an important role in normal development by regulating cell proliferation and ECM matrix remodeling [13–16], elevated expression of DDR2 has been shown to correlate with increased metastasis in various cancers [16–20].

During metastasis, cancer cells detach from the primary tumor and migrate through the ECM to invade the surrounding tissues [21]. Knocking down DDR2 has resulted in decreased cell migration [19, 22–26] in various cell types. Cell migration is regulated by multiple factors including cell-ECM adhesion, cellular contractility, and cell mechanics. However, the mechanisms by which DDR2 modulate neuroblastoma cell mechanics and cell migration are still unknown.

In the current work, we studied the cell mechanics and migration of a neuroblastoma cell line SH-SY5Y. This cell line has been extensively used as a model to study cancer progression [27]. To gain insights into the role of DDR2 and collagen binding on neuroblastoma cellular mechanics, tetracycline inducible-shDDR2 (shDDR2) was stably transfected into the SH-SY5Y cells. The utilization and engineering of shDDR2 allows for precise control of DDR2 down-regulation within these cells. Additionally, tet-inducible systems exhibit low basal expression in the absence of doxycycline, resulting in minimal interference with normal cellular processes. Together, this shDDR2 cell line allows for a precisely controlled environment to investigate the consequences of the gene manipulation under conditions that mimic disease states. Novel biophysical methods were employed to elucidate the mechanical changes from DDR2 down-regulated neuroblastoma cells. To investigate cell microrheological properties, Atomic Force Microscopy was utilized, allowing us to extract single cell stiffness measurements. Additionally, it is imperative to unveil the traction forces that shDDR2 cells exhibit in biologically relevant elastic substrates. To mimic the in-vivo environment, neuroblastoma cells were plated onto soft collagen coated elastic substrates. This allowed us to conclude changes to the shDDR2 traction forces in a biologically relevant way. To the best of our knowledge, this is the first study to unveil cellular migration mechanics on SH-SY5Y cells utilizing the novel shDDR2 cell line. Through these techniques, our results suggested increased traction forces, cell elongation, and cell stiffness were key components to mediate neuroblastoma cellular mechanics. We confirmed that on-target inhibition of DDR2 decreased its specificity for collagen, particularly regarding cellular traction forces and traction stress. DDR2 is a strong prognostic marker of poor survival in various cancers and adult tumors [28]. This work defines a novel role for this mesenchymal marker in regulating the mechanisms of neuroblastoma migration mechanics.

## 2. Materials and Methods

### Cell Culture

Human neuroblastoma cell line SH-SY5Y stably transduced with the Tet-pLKO-puro lentiviral vector expressing either a control non-targeting shRNA or shDDR2 were kindly provided by Dr. Jason Shohet (UMass Chan Medical School). Cells were cultured in Dulbecco’s modified Eagle’s medium (DMEM) supplemented with 10% fetal bovine serum (Gibco), 2mM glutamine (Gibco) and antibiotics (penicillin and streptomycin) (Gibco).

### Lentivirus Preparation and Infection

HEK-293T cells were maintained at 37°C in Dulbecco’s modified Eagle medium (DMEM), supplemented with 10% FCS and antibiotics (100 units/ml penicillin and 100 μg/ml streptomycin). Cells were transfected with pVSV-G [29] and pCMVΔR8.91 [30], together with the pLKO.1-puro non-targeting vector (Sigma Mission clone SHC016; ‘shCTRL’) or the Tet-pLKO.1-shRNA vector (Sigma Mission TRCN0000001418; ‘shDDR2’) using Lipofectamine^TM^ 2000 reagent (Thermo Fisher Scientific) as recommended by the manufacturer and following the recommendations of the RNAi Consortium (TRC) laboratory protocols with slight modifications. Twelve hours after transfection the medium was replaced by DMEM, supplemented with 30% FCS and antibiotics. Cell supernatants were harvested every 24 hours, replaced with fresh medium, and stored at 4°C until collection of the last harvest (72 hour). At this point, the consecutive harvests were pooled, filtered through 0.45mm filters and split in 3-5 mL aliquots, which were stored at −80°C. SH-SY5Y cells were infected with shCTRL or shDDR2 lentiviral particles by adding a 1:1 mix of medium:viral supernatant for 24-48 hours. Puromycin selection (2 μg/ml) was applied for 2-3 days and always compared to non-transduced control cells, which generally died within the first 24 hours.

### Western Blot

Western blot analysis was conducted using a standard protocol [31]. Briefly, cells grown to a 60-80% confluency were lysed in radioimmunoprecipitation assay (RIPA) lysis buffer (Prometheus Protein Biology Products #18-416) supplemented with Protease and Phosphatase Inhibitor Cocktails (Pierce, Thermo Scientific A32955 and A32957). Lysates were sonicated on ice, centrifuged at 15,000×g at 4 °C for 20 minutes and the soluble protein fraction was collected. Protein extracts were quantified using a Pierce BCA Protein Assay Kit (Thermo Scientific #23227). A total of 30-50 μg of proteins were separated via SDS-PAGE using Novex™ WedgeWell™ 4-20%, Tris-Glycine Mini Protein Gels (Invitrogen, Thermo Scientific XP04202BOX) and blotted onto a PVDF membrane using an iBlot transfer system and transfer stacks (Invitrogen, Thermo Scientific IB401001). Proteins were detected using SuperSignal™ West Pico PLUS Chemiluminescent Substrate (Thermo Scientific 34580). A ChemiDoc MP Imaging System (Bio-Rad) was used for chemiluminescent detection and analysis. Primary antibodies were: from Cell Signaling Technology DDR2 (#12133); from MilliporeSigma Anti-GAPDH antibody (MAB374); and from R&D Systems human phopho-DDR1/DDR2 (Y796/Y740) antibody (MAB25382).

### Lentivector Cell Maintenance

To maintain lentivector transfected cells, shCTRL cells are incubated at 37°C and 5% CO_2_ within tissue culture treated petri dishes or T-75 flasks. Cells are maintained in DMEM with 10% fetal bovine serum (Gibco), 2mM glutamine (Gibco) and antibiotics (penicillin and streptomycin) (Gibco). To maintain lentivector selection within shCTRL cells, puromycin is maintained within the culture medium at a concentration of 2 μg/ml. Medium and puromycin is refreshed every 2-3 days. Puromycin is used as a selection marker within the lentivector construct. Only cells that are integrated with the lentiviral vector construct (puromycin resistant gene) will survive puromycin selection. To maintain lentivector selection within shDDR2 cells, puromycin is maintained within the culture medium at a concentration of 2 μg/ml and doxycycline is maintained within the culture medium at a concentration of 1 μg/ml Doxycycline allows control over the Tet-On system. Without doxycycline, the gene of interest remains inactive. Upon doxycycline addition, it binds to the transactivator protein, causing a conformational change that allows the transactivator protein to bind to the tet operator sequence. Upon this change, the transcription is activated. Doxycycline and puromycin are refreshed every 2-3 days. Cells are harvested for experiments were used under passage 14.

### Collagen-Coated Glass Slides

25mm x 25 mm coverslips were placed into a coverslip holder, submerged in 70% ethanol, and sonicated for 15 minutes. The cleaned coverslips were then rinsed with DPBS and incubated in 0.1 mg/ml of Rat Tail Collagen I solution (Corning) for 1 hour at 37°C. Following the incubation, the coverslips were rinsed 3x with DPBS waiting 10 minutes per wash.

### Cell Spreading Area Measurements

Individual cells were allowed to attach onto collagen glass substrate for 30 minutes and cells were imaged using Olympus IX3 inverted microscope equipped with 40x 0.6 NA objective using the phase contrast mode. Cell area, aspect ratio, and circularity were measured using free-hand tool and shape descriptors within ImageJ (NIH). Area is defined as the area of the selection in square units. Aspect ratio is defined as the ratio of the particles fitting ellipse as a ratio of major axis divided by minor axis. Circularity is defined as the deviation of the area of a circle, with a perfect circle taken as a circularity of 1.0. Dimethyl sulfoxide (DMSO) (Sigma) was introduced to cells before plating on collagen coated glass substrates.

### Kymograph

Cells were plated onto collagen glass slides and allowed to attach for 30 minutes. After 30 minutes, cells were imaged using Olympus IX3 inverted microscope equipped with 40x 0.6 NA objective using the phase contrast mode for 30 minutes (5 second intervals). 3 cells were imaged per dish, for a total of 1.5 hours of imaging. Time-lapse images were analyzed using straight line tool in ImageJ (NIH) and the Multi Kymograph plug-in.

### Atomic Force Microscopy

All measurements were performed utilizing an MFP-3D-BIO atomic force microscope (Asylum Research, Santa Barbara, CA, USA) and DNP-10 cantilevers (Bruker, Camarillo, CA, USA) with nominal spring constant 0.06 N/m. To ensure reliability of measurements, each cantilever was calibrated for spring constant (k).

Live cells were measured 24 h after seeding on the desired surfaces to ensure proper spreading, while preventing high confluency. Measurements were limited to isolated cells to reduce the influence of cell-cell communication. Phase contrast microscopy was used in unison with AFM to align the cantilever tip over desired measurement areas of the cell. Individual force curves were taken in three locations in the perinuclear region of each cell to guarantee the thickness of the cell was significantly greater than the distance the cantilever indented into the cell. Each force curve was taken at a velocity of 2 µm/s and to a trigger point of 0.2~1 nN. 10~16 cells were measured per dish and no longer than 30 min after the dish was removed from the incubator to ensure cell viability.

Using a custom MATLAB code, cantilever deflection as a function of sample indentation depth was extracted from AFM force curves. Stiffness values were determined from the deflection-indentation curve using the Hertz model with a conical tip:

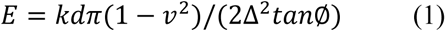

where *k* is the cantilever spring constant, *d* is the cantilever deflection, *ν* is the Poisson’s ratio value (using 0.5), Δ is the sample indentation depth, and *ϕ* is half the conical opening angle of the AFM tip [32]. To minimize the effects of nonlinear effects, force-indentation curves were fit to the Hertz model over the first 200~500 nm indentation depth.

### Polyacrylamide Substrate Preparation

Polyacrylamide gel substrates were prepared through the polymerization of 8% acrylamide and 0.04% bis-acrylamide to achieve a stiffness of 2 kPa. This polymerization process was initiated by a solution containing 0.1% ammonium persulfate and 0.3% N,N,N’,N’-tetramethylethylenediamine. 0.1 mg/mL of Collagen type I (Corning) or 10 *μ*g/mL of Fibronectin (AdvancedBioMatrix) was crosslinked to the PAA gel surface using sulfo-SANPAH. The gels were submerged under 1mg/ml sulfo-SANPAH (G-Biosciences, St. Louis, MO) solution and placed 2 inches below an 8 W ultraviolet UV lamp (Hitachi F8T5 – 365nm) and irradiated for 15 minutes. The gels were then washed with HEPES buffer and soaked with 0.1 mg/ml rat-tail collagen type I or 10 µg/mL fibronectin for 12 hours at 4 °C. After collagen coating, the collagen was aspirated, and gels were placed in culture medium and incubated for 30 minutes at 37°C before cells were seeded on them.

For traction force microscopy, we followed an established protocol [33] to fabricate gel disks of 18 mm in diameter and approximately 100 μm in thickness. These gel disks were prepared with 0.1 μm red fluorescent beads (Life Technologies) embedded just beneath the top surface. 25 mm x 25 mm square glass coverslips were cleaned, treated with 1% 3-aminopropyl-trimethoxysilane solution for 10 minutes, and then coated with 0.5% glutaraldehyde. Round glass coverslips, 18 mm in diameter, were plasma cleaned and coated with a thin layer of fluorescent beads. A 25 μL mixture of acrylamide, bis-acrylamide, and initiators was applied between the glutaraldehyde-coated square coverslip and the beads-coated round coverslip, followed by polymerization at room temperature for 15 minutes. Subsequently, the round coverslip was gently removed, leaving the resulting gel disk firmly attached to the square coverslip, with the embedded beads positioned within 2 μm below the gel surface.

### Cell Traction Force Measurements

Cell traction forces were measured using traction force microscopy [33]. Cells were cultured on PAA substrates for 24 hours before subjected to traction force microscopy. For each cell selected for traction force microscopy, a fluorescence image of the substrate was recorded to capture the marker beads in the stressed state. In addition, a phase-contrast image was acquired to record the morphology of the cell. Trypsin (Invitrogen) was then applied to disrupt cell-substrate interactions and cause the cell to detach from the substrate. A final fluorescent image of the substrate was taken to capture the marker beads in the relaxed, unstressed state. Bead displacements were calculated from the two fluorescent images using a particle image velocimetry toolbox written in MATLAB [34]. Traction stress on the gel surface was calculated from the bead displacements using the finite element analysis software (Ansys, Inc). The magnitude of total traction force (F) was calculated by integrating the magnitude of traction stress over the cell area.

### 2D Cell Migration Assay

Cells were cultured in full complete DMEM (Gibco) with the addition of HEPES to allow CO_2_ exchange for prolonged live-cell imaging. Individual cells were plated onto collagen coated slides and incubated overnight. Following incubation, cells were imaged using an Olympus IX83 inverted microscopy equipped with a 10x 0.3 NA objective using the phase contrast mode. Time-lapse images were acquired at interval of Δ*t* = 5 minutes for 4 hours in duration. Centroids of cells were analyzed from the time-lapse images using ImageJ. The mean square displacement (MSD) of each individual cell was then calculated from the cell trajectory using the MSDAnalyzer [35] for MATLAB. The MSD for each cell type was calculated from the average MSD of individual cells and was fit to the anomalous diffusion model:

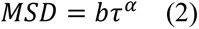

using MATLAB Curve Fitting Toolbox with *τ* representing the time interval and *α* representing the anomalous exponent. The *α*-value is an important index for directional persistence: it will be equal to 1 for cells doing normal Brownian motion and 2 for cells that move in a perfectly straight manner. The average speed of each cell was calculated from the cell trajectory by averaging the displacement magnitudes between two consecutive frames divided by Δ*t*.

### Statistics

Prism 9.0 GraphPad software was used for graph generation and statistical analyses. Significance level was set to *p* < 0.05. Number of independent biological replicates, sample sizes analyzed, and statistical tests used are stated in the figure legends.

## 3. Results

### 3.1 shRNA-mediated knockdown of DDR2

Neurobalstoma cell lines and tumors express relatively high levels of DDR2, which is significantly correlated to a worse patient survival probability [20]. Figure 1A shows the expression DDR2 mRNA across hundreds of cancer cell lines (and non-transformed control cell lines) from the Dependency Map Portal (depmap.org, Broad Institute). Cell lines are grouped by (tissue of origin/Primary tumor). As it can be clearly seen from this graph, Neuroblastoma cell lines –including SH-SY5Y cells– express a high level of DDR2 mRNA relative to most other cell lines in the study, which is evident both as the average/median expression, and for each single Peripheral Nervous System cell line (except for one cell line, which indeed corresponds to the Nerve Sheath Tumor cell line HSSCH2, not neuroblastoma). The Tet-pLKO-puro shRNA system was utilized to generate shDDR2 cells, in which DDR2 knockdown can be induced by doxycycline. Western blot analysis confirmed the efficient knock-down of DDR2 expression with doxycycline concentration ranging from 0.02 – 12.5 mg/mL (Fig. 1B). High expression level of DDR2 from SH-SY5Y was confirmed by Western blot analysis in the Tet-shDDR2-expressing SH-SY5Ycell system in the absence of doxycycline (Fig. 1B). Doxycycline treatment efficiently induced DDR2 downregulation in Tet-shDDR2 SH-SY5Y cells (Fig. 1B). Importantly, loss of DDR2 expression did not result in any gross alterations on cellular viability, except at the highest dose tested (12.5 mg/ml), an effect likely due to unspecific toxicity (Fig. 1C, S1). As expected, doxycycline-mediated downregulation of DDR2 resulted in a critical impairment of collagen-dependent autophosphorylation on tyrosine 740 of DDR2, the first and indispensable signaling event triggered upon collagen/DDR2 ligation (Fig. 1D). In conclusion, lentiviral transduction and selection were successfully achieved to make a stable SH-SY5Y cells with an inducible shRNA knockdown, with dramatic consequences in DDR2-mediated signaling.

**Figure 1.**
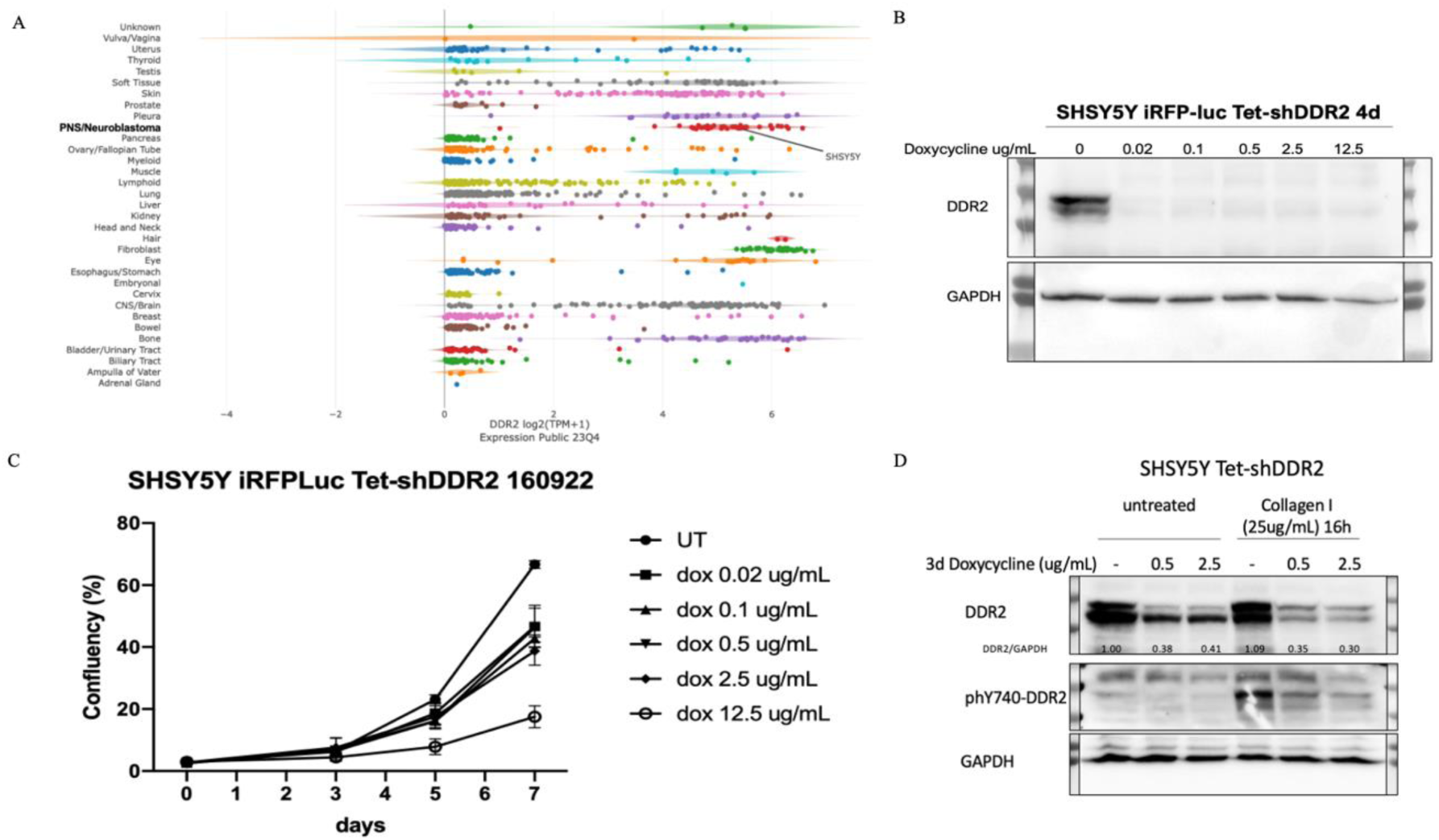
shRNA mediated knockdown of DDR2. A) DDR2 mRNA expression across cancer lines was analyzed in the Dependency Map Portal (https://depmap.org/portal; Broad Institute). B) Western blot of DDR2 expression (vs. GAPDH) upon 4 days of doxycycline treatment (0-12.5 mg/mL) on Tet-shDDR2 SHSY5Y cells. C) Cell survival/proliferation time course assay (as measured by % confluency) of Tet-shDDR2 SHSY5Y cells treated with doxycycline (on day 0) at increasing concentrations as indicated (n = 6 wells/condition). D) Western blot of DDR2 and phospho-Tyrosine 720-DDR2 (phY720-DDR2) from Tet-shDDR2 SHSY5Y cells pretreated for 3 days with the indicated concentrations of doxycycline and left unstimulated or induced with 25 mg/mL of rat tail Type I collagen for 16 hours.

### 3.2 Dependence of human neuroblastoma phenotype on collagen I and DDR2

DDR2 alterations (overexpression, amplification, and mutations) are known to drive more aggressive phenotypes in several cancer types [36–41]. Morphological adaptations and their maintenance are a consequence of the microenvironmental influence, where a cascade of interactions can determine the cellular behaviors via signaling pathway activation and cytoskeleton arrangement [42]. Here, we measured the morphological characteristics of DDR2-expressing cells and DDR2-suppressed cells. Sitravatinib, a multi-kinase inhibitor with an affinity for DDR2 and other similar kinases, has shown effectiveness in preclinical models of high-risk neuroblastoma [43]. This drug is in phase III clinical trials as it can shift the tumor microenvironment (TME) towards an immunostimulatory state [44]. When cultured on collagen I-treated glass, shCTRL cells displayed increased areas and aspect ratios with decreased circularity compared to shDDR2 and Sitravatinib-treated cells (Fig. 2, S2). Due to an increase in area and aspect ratio, shCTRL cells exhibit a more invasive, spindle-shaped cell morphology. Higher DDR2 expression levels in shCTRL cells suggest that DDR2 plays a role in maintaining an aggressive phenotype.

**Figure 2.**
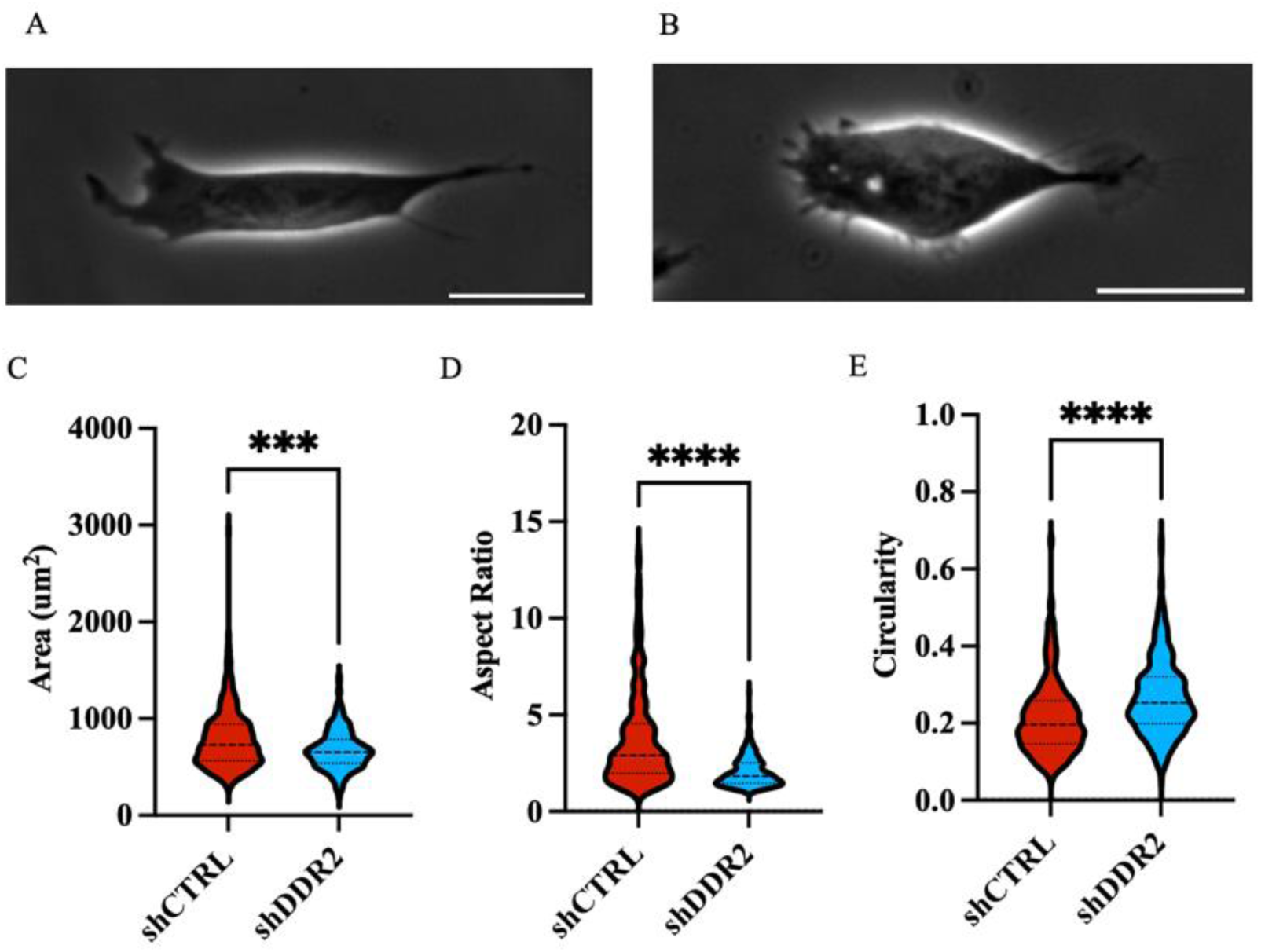
Cell morphology analysis on collagen coated glass substrates. Representative phase-contrast images of (A) shCTRL and (B) shDDR2 cells on collagen coated glass substrates. Quantification of (C) cell area (D) aspect ratio and (E) circularity. Experiments performed in three independent experiments. Un-paired t-test, p < 0.05 (n =228-329 cells). Data are presented as ± s.e.m. Scale bars represent 20 *μ*m.

### 3.3 DDR2 silencing altered cell migration on 2D collagen substrates

Mean squared displacement (MSD) is often used in physics as a metric to quantify the movement of particles. The MSD represents an average squared displacement over increasing time intervals between positions of a migration trajectory. In relation to cell migration, MSD can represent the surface area explored by the cells over a time interval, which is related to the overall efficiency of migration. Cell centroids on 2D collagen-coated glass slides were tracked for 4-hour time lapses and trajectories were plotted (Fig. 3A). Furthermore, a logarithmic scale of MSD was plotted for both shCTRL and shDDR2, and the exponent, or slope, *α*, was found to describe the migration patterns (Fig. 3B). An alpha value of 0.88 describes the shCTRL data and 1.25 describes the shDDR2 data. It is important to note that at shorter time scales shCTRL cells start in the sub-diffusive region (a = 0.89) and increases to diffusion (a = 1.02) after 150 minutes. However, shDDR2 begins in normal diffusion (a = 1.12) and switches to super-diffusive (a = 1.33) after 75 minutes. Initial dynamics of 2D cell migration typically correspond to their initial phase of particle motion. This early phase can provide insights into how cells are influenced by their intial conditions. Later time points capture the long-term average behavior of the cells. In this study, shDDR2 cells exhibited higher diffusion coefficients than shCTRl cells at both short and long time scales. To understand the overall trend that each cell line employs through 2D cell migration, the average of the MSD was plotted with standard deviations. Together, these results reveal shDDR2 exhibited higher variability, resulting in a less predictable or definative path for these cells. Simiarly, the average speed for shDDR2 cells (0.56 *μ*m/min) is greater than (p = 0.053) that of the shCTRL cells (0.44 *μ*m/min) (Fig. 3C). Together, this 2D migration study suggests that down-regulating DDR2 led to an increase in the migration of cells on 2D collagen coated surfaces.

**Figure 3.**
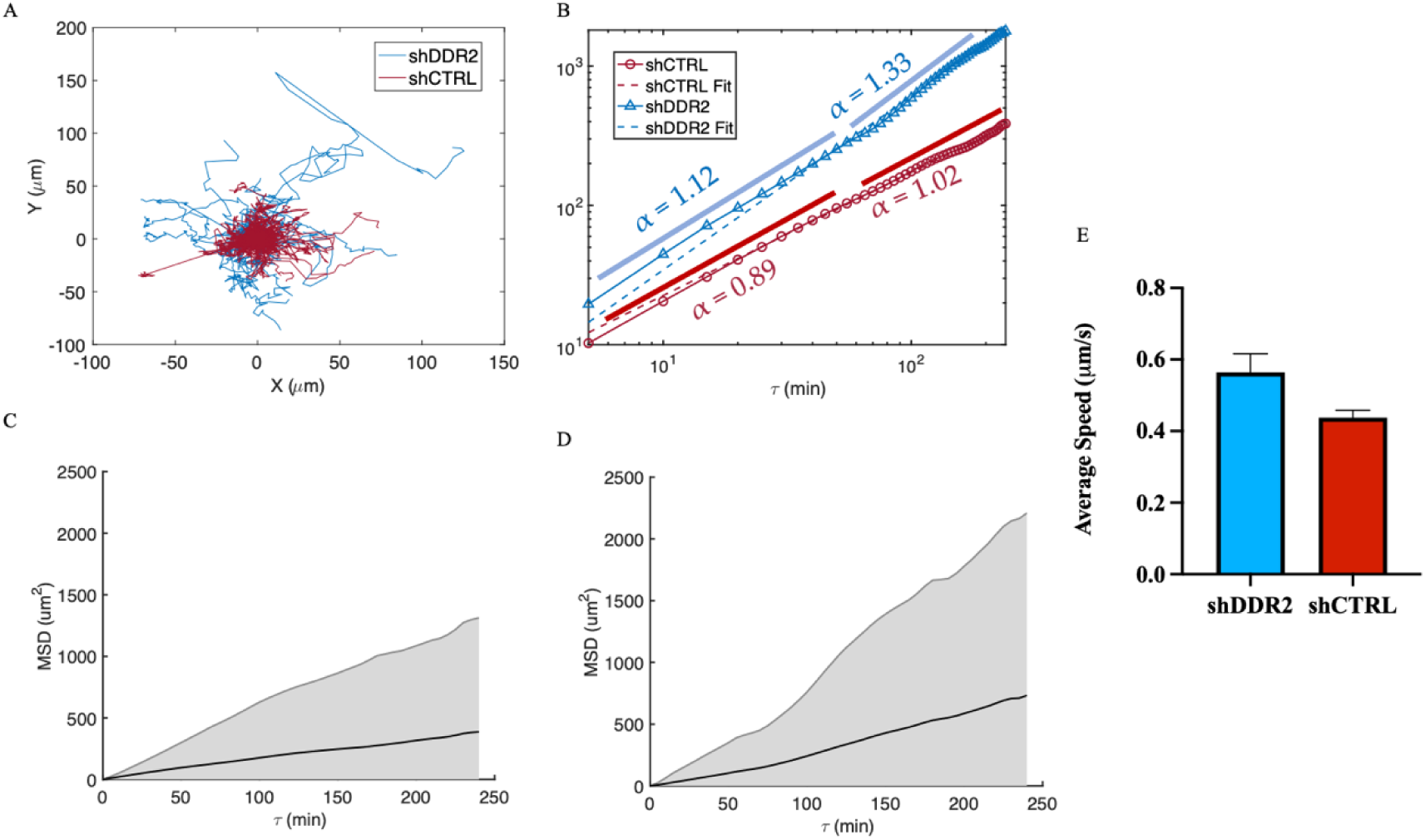
Mean Square Displacement of 2D single cell migration. (A) Individual trajectory maps of shCTRL (red) and shDDR2 (blue). (B) Log-log plot of MSD potted as a function of time interval for both shCTRL (red, α = 0.8838) and shDDR2 (blue, α = 1.247). Mean of the mean square displacement with shading representing the weighted standard deviation over all MSD for (C) shCTRL and (D) shDDR2 2D migrating cells. (E) Velocities from MSD of shCTRL and shDDR2 (n = 122 cells shCTRL, 55 cells shDDR2). Mann-Whitney Test, P = 0.0531. Data are represented as ± s.e.m.

### 3.4 DDR2 affects cellular protrusion dynamics

Cell migration involves a cyclic coordination of protrusion at the cell front, adhesion of the newly protruded domain to the substrate, and pulling of the bulk of the cell towards new adhesion sites and breaking of adhesion and retraction at the cell rear. For migration, cells must elongate and retract. Filopodia, a subcellular structure at the cell front, plays a key role in the spreading and migration of cells. Thus, changes in filopodia length during the elongation and retraction stage are crucial for effective cell migration [45].

In this work, we analyzed the front protrusion and back contraction lengths for shCTRL and shDDR2 cells. Kymographs generated from 30-minute time-lapse videos on collagen-coated glass allowed us to extract these lengths (Fig. 4A, B, C, D). The analysis revealed an increase in the front protrusion length of shCTRL and in the back contraction length compared to DDR2 downregulated cells (Fig. 4E, F). These findings suggest that DDR2 suppression leads to decreased filopodia front protrusion and rear contraction elongation. Overall, our results indicate that DDR2 expression may promote cell mechanosensing and migration in vitro through filopodia extension.

**Figure 4.**
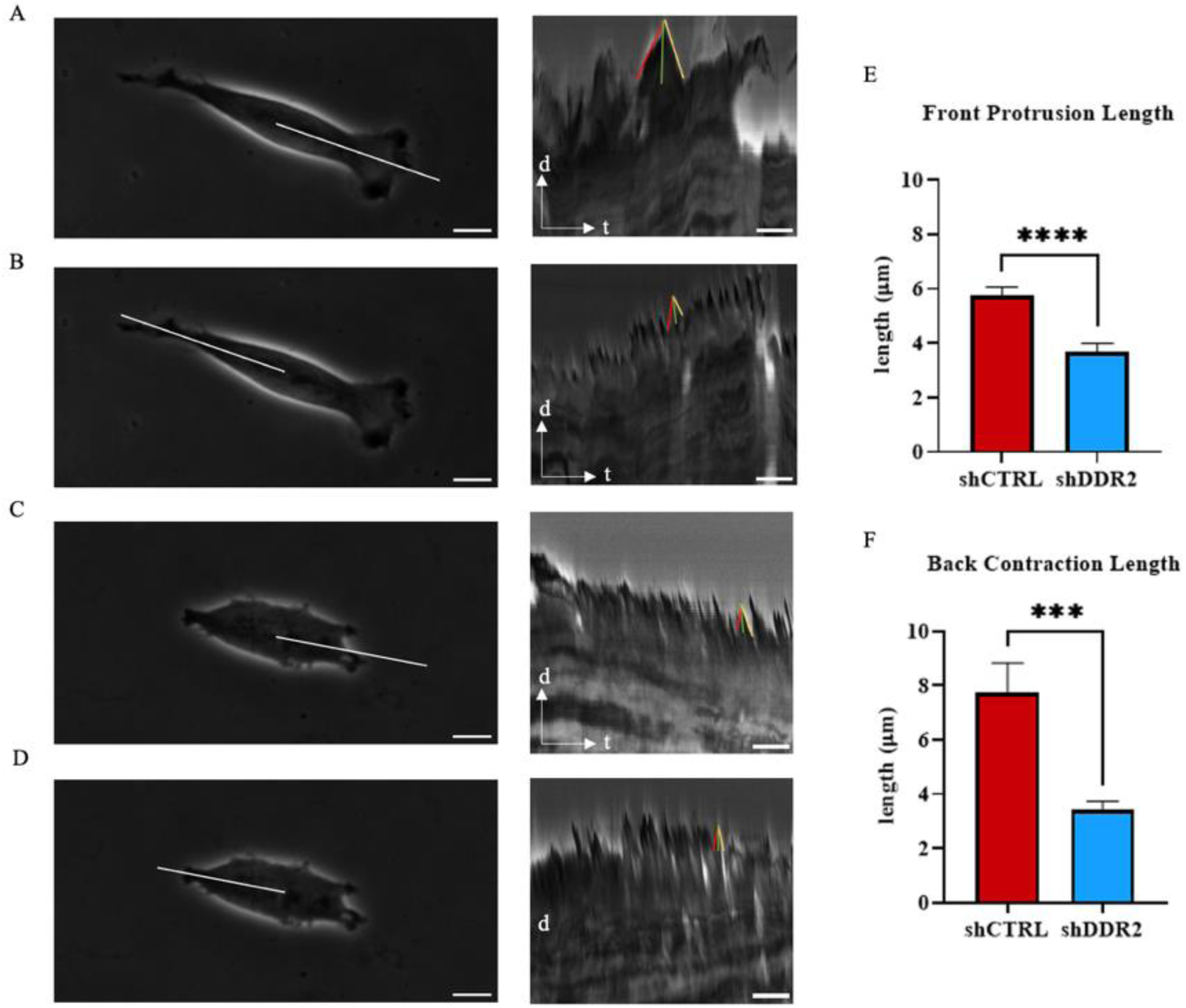
Protrusion lengths of cells on collagen coated substrates. Representative phase-contrast image of cells on collagen coated glass substrate with kymographic analysis of white line on shCTRL cell (A) front and (B) back and shDDR2 cells (C) front and (D) back. Kymograph derived (E) front protrusive length and (F) back contraction length. Red line represents protrusion, yellow line represents contraction, and green line represents event length. Experiments performed in three independent experiments. Mann-Whitney test, p < 0.05. Data are presented as ± s.e.m. Scale bars on left images represent 10 *μ*m (A-C) and in kymographs represent 250 seconds (B-D).

### 3.5 Regulation of cell stiffness with DDR2 collagen activation

Cell stiffnesses can be related to the cytoskeletal structures within the cells where low cell stiffness is attributed to disorganization to cytoskeletal structures and reduction of actin filaments [46]. To analyze the relationship between cell stiffness and DDR2 expression, we investigated the local cell stiffness using Atomic Force Microscopy (AFM). Measurements were taken at the front, middle, and rear of cells cultured on collagen-coated glass. For the detection of the Young’s modulus, the AFM cantilever tip is pressed onto the cell surface, with the young’s modulus defined as the ratio between the measured stress and strain. Utilizing AFM enabled us to pinpoint cell stiffnesses at the cell front, back, and the center (Fig. 5A). A comparison between the front and middle parts of the cell revealed significantly higher stiffnesses for shCTRL (Fig. 5B,C). However, no difference was observed in the rear stiffness between shCTRL and shDDR2 cells (Fig. 5D). These results indicate that downregulating DDR2 in neruobastoma cells decreases the cell stiffness.

**Figure 5.**
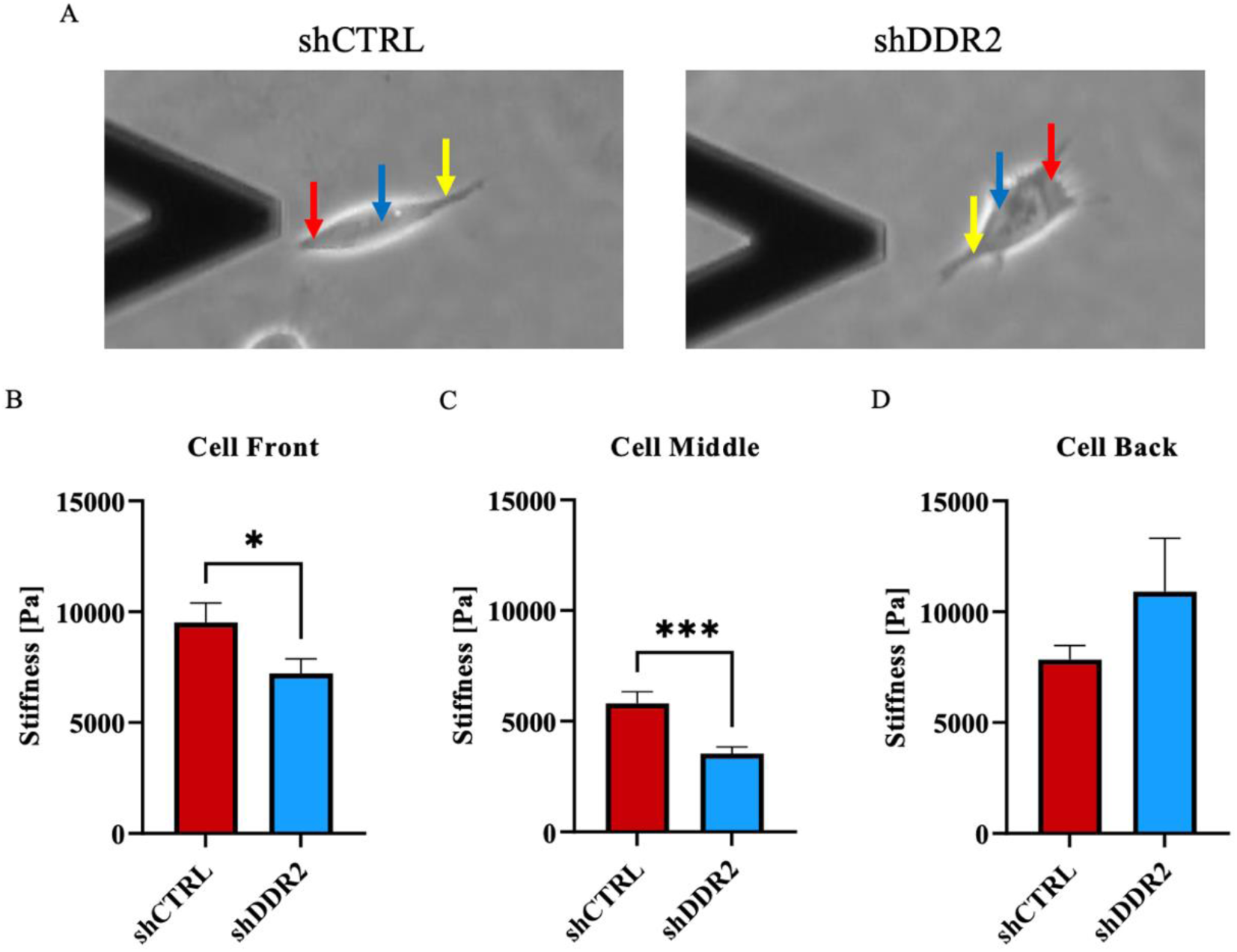
DDR2 downregulation decreases neuroblastoma cell stiffness. Youngs modulus measurements of shCTRL and shDDR2 cells using atomic force microscopy. Indentation of cell front (red arrow), cell middle (blue arrow) and cell back (yellow arrow). Young’s modulus of Experiments performed in three independent experiments. Un-paired t-test, p < 0.05 (n= 32-33 cells). Data are presented as ± s.e.m.

### 3.6 Effect of DDR2 knockdown on cell traction force

To detect and interpret the biological information in the ECM, cells adhere and transduce myosin-generated traction forces through integrin-based adhesions. This process, which provokes dynamic signaling events, is termed mechanotransduction [47]. A reduction in net adhesion and migration rates, attributed to decreased binding strength and force generation at the leading edge, can lead to partial or complete loss of migration [48]. To understand the consequences of DDR2 down-regulation on mechanotransduction, traction force measurements were taken on 2kPa PAA gels for both shCTRL and shDDR2 cells, replicating the stiffness of the human brain (Fig. 6A, B, C, D, E, F). Cellular traction forces have been shown to mediate mechanotransduction, cell migration, adhesion, and ECM remodeling [49]. shDDR2 cells exhibited dramatically reduced traction force compared to shCTRL. A significance difference was observed in the percentage of cells attached to the PAA gels with and without collagen for both shCTRL and shDDR2 cells (Fig. S3). The absence of collagen I crosslinked onto PAA gels resulted in decreased stress and total force in shCTRL cells (Fig. 6G, H). To contrast DDR2 and Collagen, traction force microscopy (TFM) was performed on fibronectin-coated surfaces for both DDR2 down-regulated cells and control cells. Contrary to collagen, in the presence of a different ECM protein, fibronectin, shCTRL and shDDR2 did not result in any differences to their traction force (Fig. S4). This data highlights the specificity of shCTRL cells for collagen. In summary, these experiments suggest that DDR2 down-regulated neuroblastoma cells exhibit altered mechanotransduction activity when plated on Collagen I cross-linked PAA gels.

**Figure 6.**
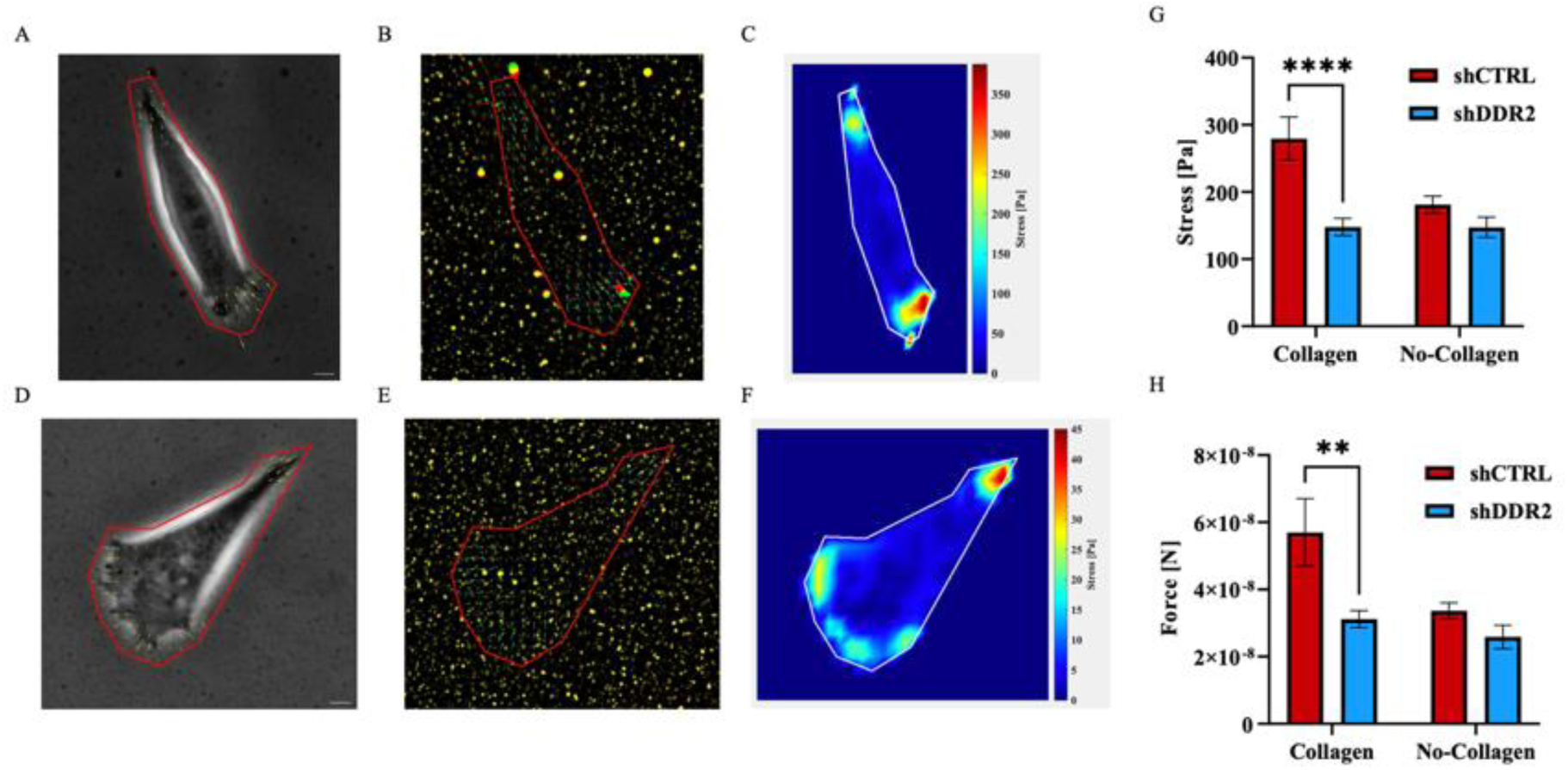
Traction force microscopy on 2kPa Collagen PAA Gels. Representative images of phase contrast, force displacement field, and stress map of shCTRL (A-C) and shDDR2 (D-F) cells. Quantification of (G) maximum traction stress and (H) total force of cells on 2kPa PAA gels (Collagen n = 60-65 cells, No-Collagen n = 45-112 cells). Experiments performed in three independent experiments. Šídák Two-Way ANOVA, p< 0.05. Data are presented as ± s.e.m. Scale bars represent 20 *μ*m.

## 4. Discussion

Cancer invasion is inherently coupled to cell mechanics and ECM properties [50–52]. These cellular mechanics, intertwined with matrix mechanics, allow cells to remodel the surrounding matrix environment by exerting forces on it. DDR2 has been shown to play a role in the growth and metastasis of epithelial and mesenchymal cancers [53–59]. However, the role DDR2 plays in the metastasis of neuroblastoma remain ill-defined.

In the present study, we provide to our knowledge the first study utilizing shRNA technology on SH-SY5Y cells to elucidate the role of DDR2 in mediating neuroblastoma cell mechanics and migration. One of the most prominent characteristics of cancer cells is their ability to proliferate constantly. We found that downregulating DDR2 was a critical regulator of neuroblastoma cell maintenance. Most receptor tyrosine kinases (RTKs) bind to soluble growth factors and mediate various cellular responses, including proliferation, differentiation, migration, and survival [60, 61]. Various studies have demonstrated that DDR2 regulates cellular proliferation in embryonic cells [62], breast cancer cells [63], endochondral cells [64], and squamous cells [65]. Therefore, it is not surprising that DDR2 is also essential for neuroblastoma growth.

Inhibiting DDR2 has been shown to decrease the migration of melanoma [23], fibroblasts [66], breast cancer [19] and lung cancer [67]. Previous migration studies on DDR2 have been conducted through long-term (6-24 hour) migration assays, by measuring the amount of cells invaded from populated regions into an previously empty region [22, 23]. However, both cell migration and cellular proliferation contribute to these cell invasion studies, making it hard to decouple the singular invasion effect. In this work, we directly measured the effects of downregulating DDR2 expression on the 2D migration of neuroblastoma by tracking individual cell‘s MSD, which has been utilized as a tool to gain insight on the migratory mode [68–70]. Notably, MSD for both shCTRL and shDDR2 exhibit a non-linear increase with τ, indicating anomalous dynamics of their migration. At shorter time scale (*τ* < 60 minutes), the migration of both cell lines can be modeled as normal Brownian motion, with anomalous exponent *α* approximately equal to 1. At longer time scale, the *α* value of shDDR2 cell line increased to 1.3, suggesting that knocking down DDR2 shifts the cell migration mode from diffusive mode to super-diffusive mode as time increases. Although our results show no statistical significance (*p* = 0.053), there is an increased trend for the average migratory speed for the DDR2-downregulated cells. The biphasic relationship between cell migration and adhesion has been well studied [71, 72]. DDR2 depletion has been shown to decrease the cell focal adhesion size [73] which consequently alters the migration of cells [74]. With the knowledge that DDR2 autophosphorylation begins after binding to collagen, it is hypothesized that this lack of DDR2 binding to collagen in shDDR2 cells causes an increase to the MSD anomalous exponent into the super-diffusive region.

shDDR2 SH-SY5Y cells also reduced their cell area and aspect ratios, consistent with other reports [22]. In addition to morphology changes, the net traction forces, frontal protrusion length and stiffnesses decreased for the DDR2-downreulated cells. These results suggest that DDR2 could modulate a mesenchymal phenotype [75]. Some reports associate decreased cell stiffness to cancer invasiveness [76, 77]. However, cancer cell invasion is a complex process that cannot be predicted by a single cell parameter alone [78]. As a first step of mesenchymal cell migration, cells develop a leading edge that mechanically couples to the extracellular tissue which is followed by cell contraction, gradual sliding of the cell rear, and translocation of the cell body [79]. The filopodia at the leading edge positively controls mesenchymal migration directionality and persistence [80, 81]. The data presented in this work shows the coupling of increasing protrusion and retraction lengths which compliments the increased aspect ratios of the cellular morphologies. Together, our data suggests that DDR2 effects the ability of these cells to elongate this filopodia and continue in metastatic cascade. Cell stiffness can often be attributed to the organization and bundling of actin filaments [82] which is likely due to the increase in protrusion lengths for the neuroblastoma cells studied here. Bayer *et al.* [73] found decreased cell spreading coupled with reduced traction forces on DDR2 depleted cancer associated fibroblasts (CAFs), which agrees with our findings in SH-SY5Y neuroblastoma cells. Additionally, they found that DDR2 was required for full activation of collagen binding to *β*1-containing Integrins. It is plausible that DDR2 modulates integrin-mediating mechanics to collagen substrates in this neuroblastoma cell line. In corroboration with this, we show that it is not migration speed alone that controls the metastatic ability of SH-SY5Y cells, but rather an interplay between the proliferation, protrusion dynamics, traction forces and cell stiffness. In summary, we provide possible mechanical changes from collagen binding receptor, DDR2, in neuroblastoma cell line, SH-SY5Y, for controlling the mesenchymal phenotype. We also show that DDR2 is a critical pathway to control aggressive behavior that forms in neuroblastoma and that facilitates tumor cell invasion. However, it is important to note that these changes may vary between cell lines. As such, DDR2 could represent an important therapeutic target carry significant impact on cancer progression. Our results also shed light into the action DDR2 plays in neuroblastoma cells which may contribute to migration mechanics and cell-cell and cell-ECM interactions.

## 5. Conclusions

In this study, we used a normal and DDR2 downregulated neuroblastoma cell line to conduct a study on cellular mechanics of DDR2 upon activation to Collagen I. We employed various parameters to verify the robustness of our results. The combined results obtained for control and DDR2 down-regulated cells suggest that collagen activation of DDR2 tyrosine kinase is required for the normal neuroblastoma growth and spreading. Similarly, neuroblastoma cells with down-regulated DDR2 exhibited significantly decreased cell stiffnesses, protrusion lengths, and consequently, lower traction forces compared to control neuroblastoma cells. However, a dynamic balance between these variables is necessary as an increase to cell stiffness causes a decrease to the MSD. These data signify the importance of DDR2 and collagen signaling by elucidating the mechanisms involved in the mesenchymal neuroblastoma cell line migration mechanics. Our results provide insight into the mechanics of collagen-binding receptor DDR2 as a critical pathway for controlling neuroblastoma cells.

## Author Contributions

Data curation, Theadora Vessella and Esteban Rozen; Formal analysis, Theadora Vessella and Esteban Rozen; Investigation, Hong Susan Zhou, Qi Wen and Jason Shohet; Resources, Hong Susan Zhou, Qi Wen, Esteban Rozen and Jason Shohet; Supervision, Hong Susan Zhou, Qi Wen and Jason Shohet; Writing – original draft, Hong Susan Zhou, Qi Wen, Theadora Vessella and Esteban Rozen; Writing – review & editing, Hong Susan Zhou, Qi Wen, Theadora Vessella, Esteban Rozen and Jason Shohet. All authors have read and agreed to the published version of the manuscript.

## Acknowledgments

The authors would like to acknowledge Sara VanHosen for help with kymograph analysis and Jiazhang Chen for help with Atomic Force Microscopy measurements.

