## Supplemental Information for "Investigation of cell mechanics and migration on DDR2-expressing neuroblastoma cell line"

Theadora Vessell$a^{1}$, Esteban Roze$n^{3,4}$, Jason Shohe$t^{4}$, Qi We$n^{2}$, Hong Susan Zho$u^{1}$

1 Department of Chemical Engineering, Worcester Polytechnic Institute, 100 Institute Rd, Worcester Massachusetts 01609, USA

2 Department of Physics, Worcester Polytechnic Institute, 100 Institute Rd, Worcester Massachusetts, 01609, USA

3 Crnic Institute Bolder Branch, BioFrontiers Institute, University of Colorado Boulder, 3415 Colorado Avenue, Boulder Colorado, 80303, USA

4 University of Massachusetts Medical School, Department of Pediatrics, 55 Lake Ave North, Worcester Massachusetts, 015656, USA

Keywords: Cancer Metastasis, DDR2, neuroblastoma

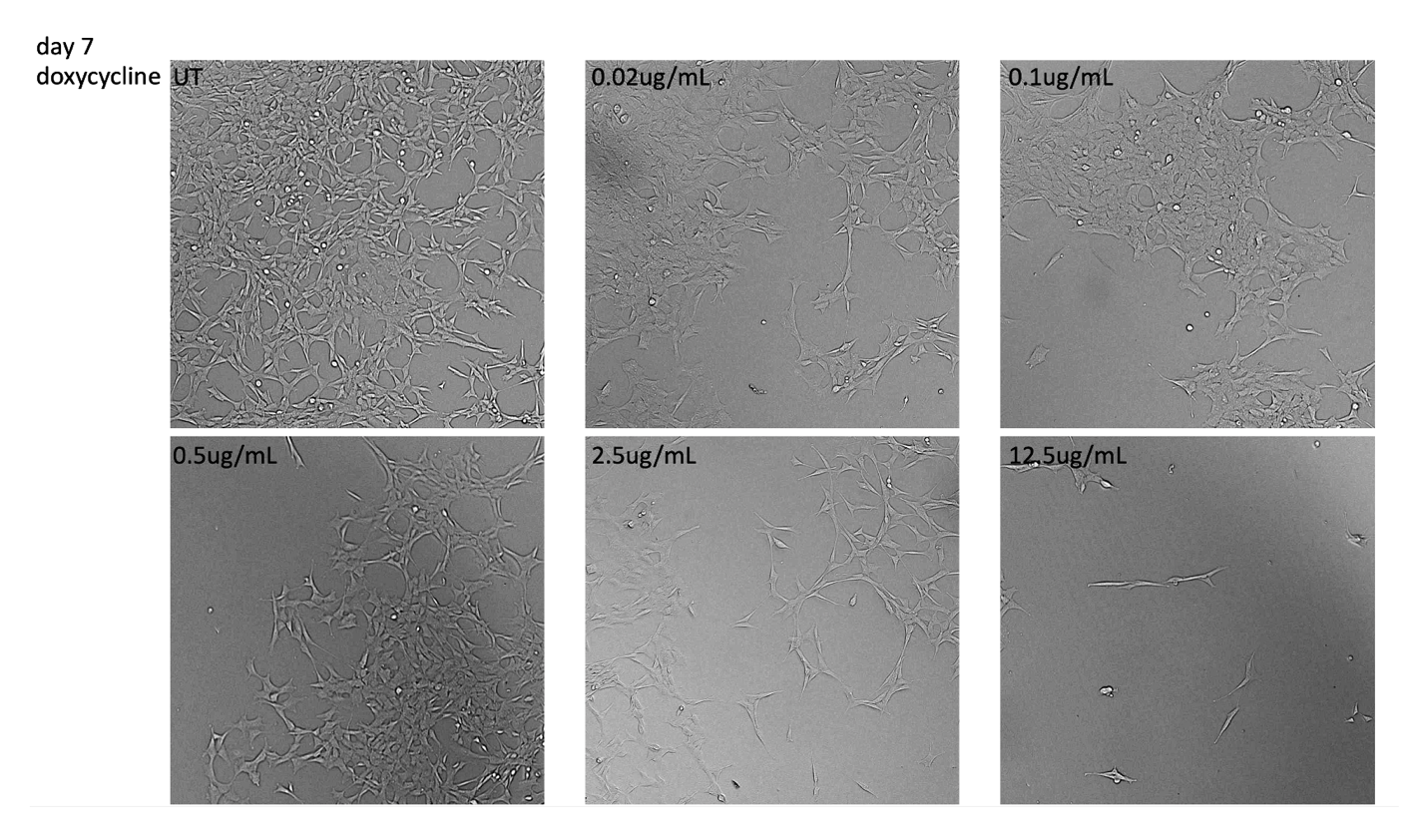

**Figure S1. shiDDR2 confluency with treatment of doxycycline.** Representative bright field images of shDDR2 cell line with untreated, 0.02 mg/mL – 15 mg/mL of doxycycline.

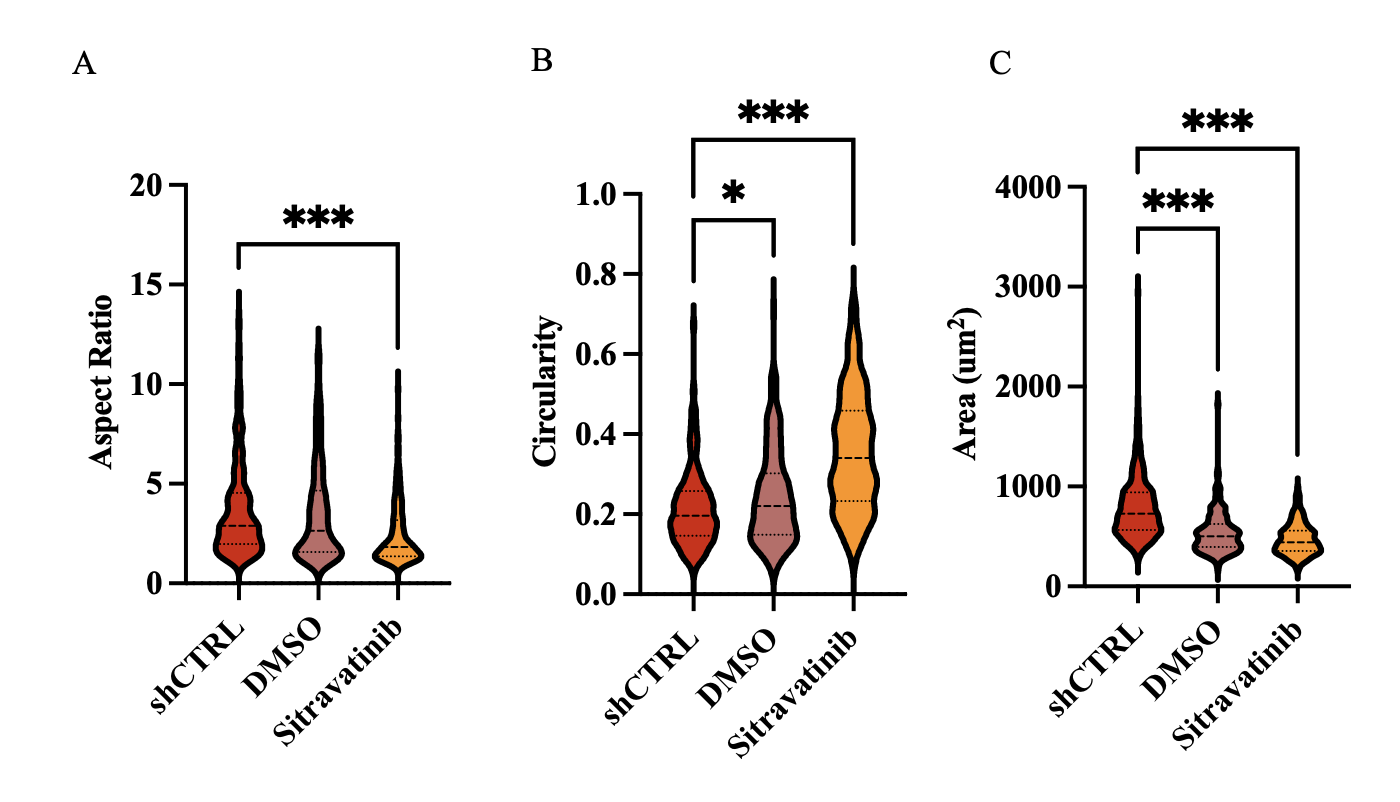

**Figure S2. Morphology of cells attached to collagen coated glass substrates morphology.** Quantification of A) cell area B) cell aspect ratio and C) cell circularity of control, control cells treated with 0.05% DMSO, and Sitravatinib treated cell. Experiments performed in three independent experiments (n=184-329 cells). Un-paired t-test, p < 0.05. Data are presented as

± s.e.m. Scale bars represent 20 𝜇m.

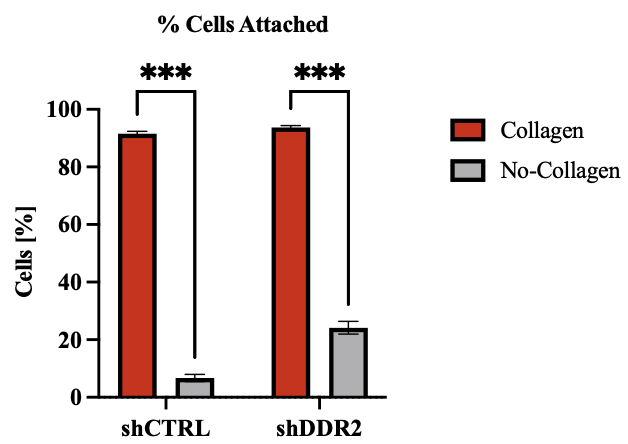

**Figure S3. Percentage of cells attached onto 2kPa PAA gel.** Percentage of cells that remain attached to collagen coated PAA gel from before and after vigorous rinsing with DPBS. Experiments performed in three independent experiments. Tukey 2Way ANOVA, P<0.05. Data are presented as

± s.e.m.

 
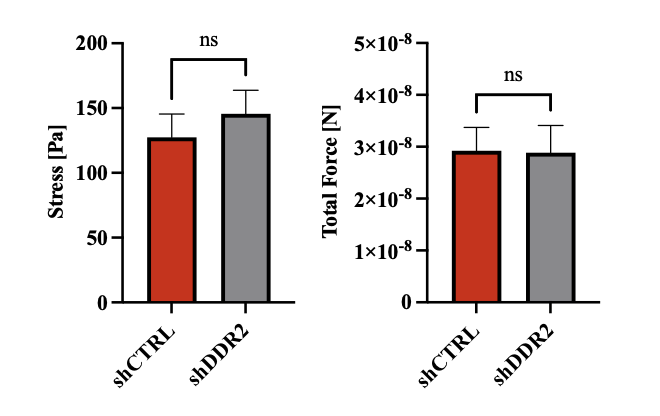

**Figure S4. Traction force microscopy on fibronectin coated 2kPa PAA Gel.** A) total force and B) maximum traction stress (n= 22 - 27 cells). Experiments performed in three independent experiments. Unpaired t-test, P<0.05. Data are presented as ± s.e.m.
